# Revealing Hi-C subcompartments by imputing high-resolution inter-chromosomal chromatin interactions

**DOI:** 10.1101/505503

**Authors:** Kyle Xiong, Jian Ma

## Abstract

The higher-order genome organization and its variation in different cellular conditions remains poorly understood. Recent high-resolution genome-wide mapping of chromatin interactions using Hi-C has revealed that chromosomes in the human genome are spatially segregated into distinct subcompartments. However, due to the requirement on sequencing coverage of the Hi-C data to define subcompartments, to date subcompartment annotation is only available in the GM12878 cell line, making it impractical to compare Hi-C subcompartment patterns across multiple cell types. Here we develop a new computational approach, named Sniper, based on an autoencoder and multilayer perceptron classifier to infer subcompartments using typical Hi-C datasets with moderate coverage. We demonstrated that Sniper can accurately reveal subcompartments based on Hi-C datasets with moderate coverage and can significantly outperform an existing method that uses numerous epigenomic datasets as input features in GM12878. We applied Sniper to eight additional cell lines to identify the variation of Hi-C subcompartments across different cell types. Sniper revealed that chromosomal regions with conserved and more dynamic subcompartment annotations across cell types have different patterns of functional genomic features. This work demonstrates that Sniper is effective in identifying subcompartments without the need of high-coverage Hi-C data and has the potential to provide new insights into the spatial genome organization variation across different cell types.

## Introduction

In humans and other higher eukaryotes, chromosomes are folded and organized in 3D space within the nucleus and different chromosomal loci interact with each other (Bickmore and van Steensel, 2013; Bonev and Cavalli, 2016; Rowley and Corces, 2018). Recent developments in whole-genome mapping of chromatin interactions, such as Hi-C (Lieberman-Aiden et al., 2009; Rao et al., 2014), have facilitated the identification of genome-wide chromatin organizations comprehensively, revealing important 3D genome features such as loops (Rao et al., 2014), topologically associating domains (TADs) (Dixon et al., 2012; Sexton et al., 2012; Nora et al., 2012), and A/B compartments (Lieberman-Aiden et al., 2009). Specifically, at megabase resolution, chromosomes are largely segregated into two compartments, A and B (Simonis et al., 2006; Lieberman-Aiden et al., 2009). Compartment A regions generally contain open and active chromatin, while compartment B regions are mostly transcriptionally repressed. Further analysis showed that these A/B compartment domains can be inferred from epigenetic status including DNA methylation and chromatin accessibility (Fortin and Hansen, 2015). The separations of B and A compartments in the genome also have near identical agreement with lamina associated domains (LADs) and inter-LADs, respectively (Kind et al., 2015; van Steensel and Belmont, 2017), suggesting that A/B compartments have different spatial positions in the nucleus. More recently, A/B compartment separations have been observed using other genomic and imaging approaches to probing the 3D genome (Quinodoz et al., 2018; Beagrie et al., 2017; Wang et al., 2016).

In Rao et al. (2014), the A/B compartment definitions were greatly enhanced using high-resolution (up to 1kb) Hi-C data generated from the human lymphoblastoid (GM12878) cell line. Specifically, Rao et al. (2014) identified Hi-C subcompartments that divide A/B compartments into five primary subcompartments: A1, A2, B1, B2, and B3. These Hi-C subcompartments show distinct and more refined associations with various genomic and epigenomic features such as gene expression, active/repressive histone marks, DNA replication timing, and specific subnuclear structures (Rao et al., 2014). A more recent study based on the new TSA-seq technology further demonstrated that these subcompartments strongly correlate with cytological distance between the chromatin and specific subnuclear structures such as nuclear speckles and nuclear lamina, reflecting the spatial localization of the chromatin in the nucleus (Chen et al., 2018). Therefore, the annotation of Hi-C subcompartments could be extremely useful to provide complementary perspective of the 3D genome in terms of its spatial position in cell nucleus.

Unfortunately, Hi-C data from GM12878, which has almost 5 billion mapped paired-end read pairs, is the only dataset with sufficient coverage to allow reliable identification of subcompartments through clustering inter-chromosomal contact matrices. When the same clustering procedure is applied on lower coverage inter-chromosomal contact maps from most available Hi-C datasets that typically have 400 million to 1 billion mapped reads (Rao et al., 2014), the inter-chromosomal contact matrices are often too sparse to reveal clear subcompartment patterns. Recently, a neural network based method called Megabase was developed to predict Hi-C subcompartment assignments of chromosome regions with 100kb resolution using numerous epigenomic signals as features without using Hi-C data (Di Pierro et al., 2017). Based on 84 protein-binding and 11 histone marks ChIP-seq datasets in GM12878, Megabase was trained to predict the original subcompartment annotations in GM12878 from Rao et al. (2014) with over 60% consistency in each subcompartment compared to the original annotations (except for the B2 subcompartment). However, most cell types do not have as many ChIP-seq datasets as GM12878 does and some histone marks may even exhibit drastically reduced abundance in other cell lines (Yan et al., 2015). Therefore Megabase has limited application to most cell types and it is also unclear how Megabase would perform in other cell types. Indeed, comparing Hi-C subcompartments across different cell types still has not been possible.

Here we develop a new computational method called Sniper, for nuclear genome subcompartment inference using imputed probabilistic expressions of high-resolution inter-chromosomal Hi-C contacts. We utilize a neural network framework based on a denoising autoencoder (Vincent et al., 2008) and multi-layer perceptron (MLP) classifier (Haykin et al., 2009) that uses modest coverage Hi-C contact maps, which are typically available, to recover high coverage inter-chromosomal contact maps and predict the subcompartment labels of genomic regions in 100kb resolution. A recently developed method HiCPlus (Zhang et al., 2018) used convolutional neural networks (Schmidhuber, 2015) to impute intra-chromosomal chromatin contacts, but as of now there are no methods to directly impute inter-chromosomal contacts. We demonstrate that Sniper can accurately recover high coverage inter-chromosomal Hi-C contact maps in GM12878 such that we can reliably annotate subcompartments, and can significantly outperform Megabase. We applied Sniper to additional eight cell lines, including K562, IMR90, HUVEC, HeLa, HMEC, HSPC, T Cells, and HAP1, to reveal Hi-C subcompartment changes across cell types for the first time. We believe that Sniper is a useful method to offer new perspectives of genome organization changes with respect to Hi-C subcompartments in different cell types. The source code of Sniper can be accessed at: https://github.com/ma-compbio/SNIPER.

## Results

### Overview of Sniper

The overall goal of Sniper is to use only moderate coverage Hi-C data (e.g., approx. 500 million mapped read-pairs) as input to infer subcompartment annotations (Fig. 1A). Rao et al. (2014) originally defined subcompartments by using the inter-chromosomal Hi-C matrix from GM12878, constructed from Hi-C contacts between odd-numbered chromosomes along the rows and even-numbered chromosomes along the columns. They used a Gaussian hidden Markov model (HMM) to cluster on the rows of the inter-chromosomal matrix. Loci in odd-numbered chromosomes were assigned to five clusters corresponding to the five primary subcompartments. Clusters were separated into A1, A2, B1, B2, or B3 subcompartments based on the Spearman correlations between clusters. To define subcompartments in even-numbered chromosomes, Rao et al. (2014) applied the clustering method to the transpose of the inter-chromosomal matrix.

**Figure 1:**
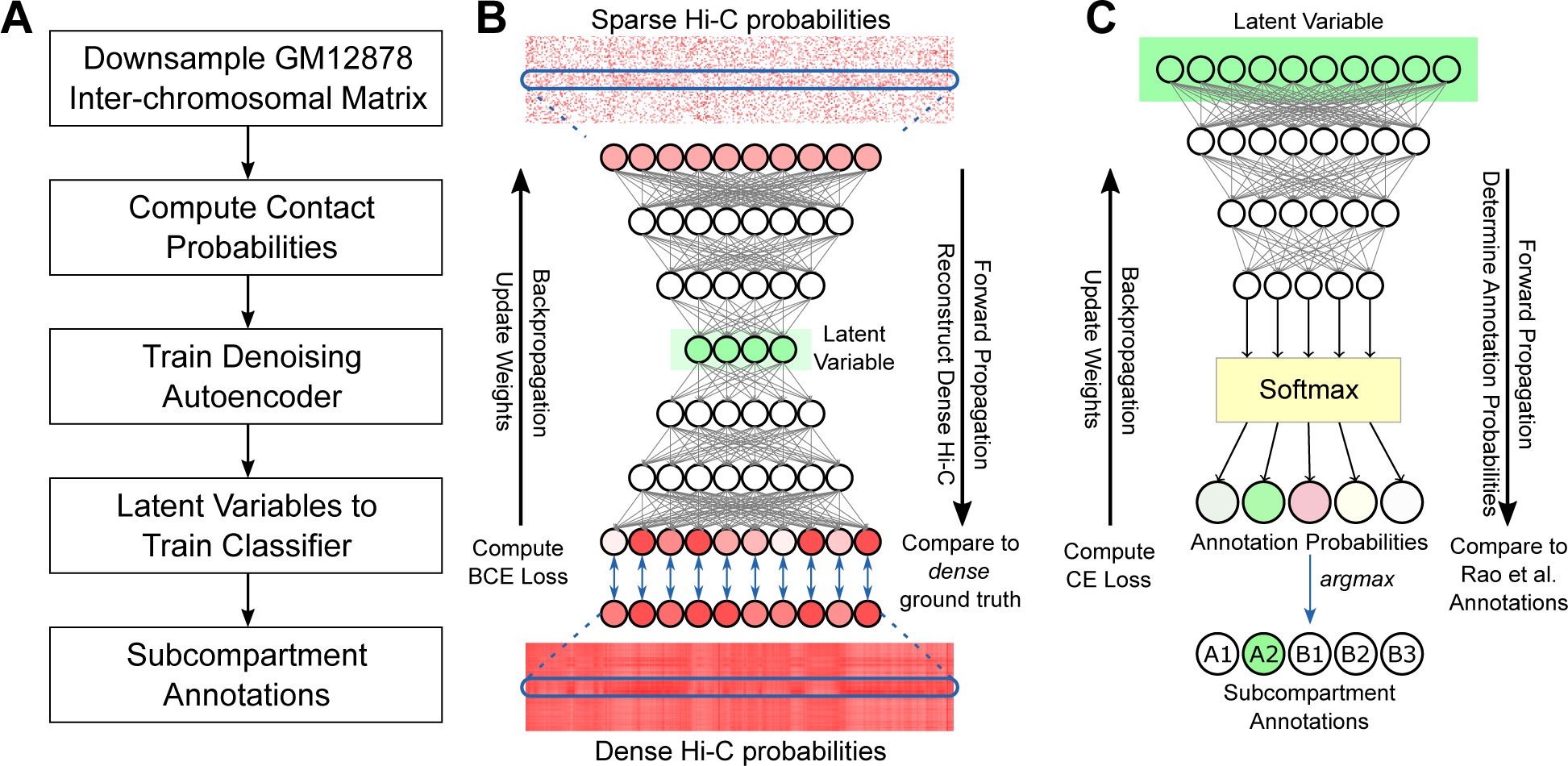
Overview of Sniper. **(A)** Flowchart of Sniper’s training procedure. **(B)** Sniper denoising autoencoder. Rows of the low coverage Hi-C probability map are used in the input layer. Weights are optimized using binary cross-entropy (BCE) loss between the reconstructed and ground truth contact probabilities. **(C)** Sniper neural network classifier is trained using latent variables from **(B)** as input and optimized using cross-entropy between predictions and the original annotations based on the high coverage Hi-C data in Rao et al. (2014).

The Sniper framework is comprised of two separate neural networks, a denoising autoencoder (Vincent et al., 2008) (Fig. 1B) and a MLP classifier (Fig. 1C). The autoencoder takes as inputs rows in a sparse inter-chromosomal Hi-C matrix (in 100kb resolution) for genomic regions in odd-numbered chromosomes along the rows and regions in even-numbered chromosomes along the columns. The autoencoder outputs dense contacts between a given region in odd-numbered chromosomes and all regions in even-numbered chromosomes. At the same time, its encoder outputs low-dimensional latent variables that represent features in the sparse matrix which capture dense chromatin contacts. The latent variable compresses high-dimensional genome-wide contacts of each genomic region into a much lower dimension, and is subsequently input into the classifier that categorizes the regions into one of five primary subcompartment classes – A1, A2, B1, B2, and B3 (based on GM12878 annotations). Note that although Rao et al. (2014) defined an additional B4 subcompartment, it is only present and specifically defined in chromosome 19, occupying less than 0.4% of the genome. We therefore did not train Sniper to consider B4. We then train a separate autoencoder and classifier to annotate regions in even-numbered chromosomes. We convert Hi-C contacts into contact probabilities to mitigate the effects of extreme Hi-C signals (see Methods). By using low dimensional representations of complex genome-wide chromatin contacts, we can predict subcompartment annotations using a basic multi-layer perceptron network. A detailed description of Sniper is provided in the Methods section.

Note that GM12878 has very high Hi-C coverage (approx. 5 billion mapped read pairs) while other cell types typically have just a few hundred million reads. To reflect coverage in other cells types, we downsampled GM12878’s Hi-C dataset to around 500 million reads by randomly removing 90% of its original reads. The inter-chromosomal Hi-C matrix from GM12878’s downsampled data is then used to train the autoencoder. Hi-C data of lower coverage cell lines can then be input into the trained networks to infer their dense Hi-C matrices and subcompartment annotations.

### Sniper can accurately predict Hi-C subcompartments in GM12878

We first evaluated the performance of Sniper in inferring subcompartments in GM12878 using downsampled Hi-C data because the annotation based on high-coverage Hi-C is readily available from Rao et al. (2014). We use confusion matrices to assess the overall accuracy of Sniper compared to Rao et al. (2014) annotations in GM12878 and also show performance differences in different subcompartments. We define accuracy as the fraction of 100kb chromatin regions whose Sniper annotations match Rao et al. (2014) annotations. The neural networks in Sniper expect inputs with the same length, but the inter-chromosomal Hi-C matrix is not symmetric. We cannot simply transpose the matrix and use a single Sniper model to predict subcompartment annotations in both odd and even-numbered chromosomes. We therefore trained two separate models of Sniper, one to classify subcompartments in odd-numbered chromosomes, and one for predictions in even-numbered chromosomes. The odd chromosome model is trained on loci from chromosomes 1, 3, 5, and 7 and tested on loci in the remaining odd-numbered chromosomes. Similarly, the even chromosome model is trained using chromosomes 2, 4, 6, 8, and 10 and tested on the remaining even-numbered chromosomes.

Sniper’s annotations of A1, A2, B1, B2, and B3 for each 100kb genomic region in GM12878 match 91.0%, 97.1%, 84.2%, 81.8%, and 93.9% of Rao et al. (2014) subcompartment annotations, respectively (Fig. 2A). Using chromosomes 9, 11, 13, 15, 17, 19, and 21 as the training set for the odd chromosome model, Sniper achieves similarly high accuracy (Fig. S1). The average precision of Sniper predictions in each subcompartment also remains high, with areas under the precision-recall curve (AUPR) of 0.981, 0.974, 0.939, 0.957, and 0.974, respectively (Fig. 2B). In 10-fold cross validation, Sniper’s accuracy remains high with low variance among training folds (Fig. 2C). Latent variables for all chromatin regions were divided into 10 partitions, each of which achieved similar accuracy compared to Rao et al. (2014) annotations (Table S1).

**Figure 2:**
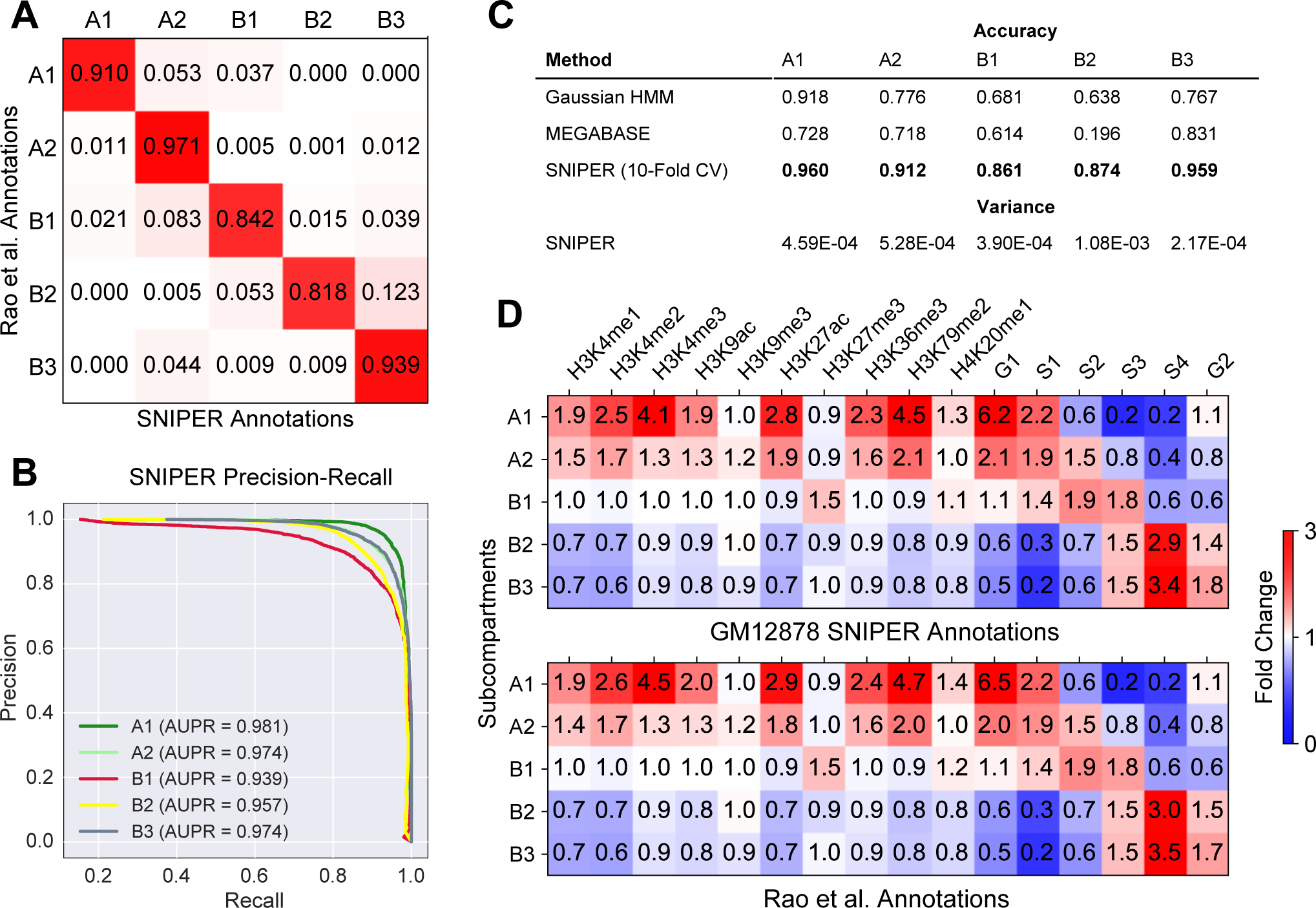
Sniper performance in GM12878. **(A)** Confusion matrix between Sniper predictions and the original subcompartment annotation based on the high-coverage Hi-C in Rao et al. (2014). **(B)** Precision-recall curve and AUPR values for the prediction of each subcompartment. **(C)** Accuracy of predicting GM12878 annotations for 100kb bins using the baseline Gaussian HMM, Megabase (Di Pierro et al., 2017), and Sniper (average across 10-fold cross validation). Highest prediction accuracy for each subcompartment is highlighted. **(D)** Histone mark and replication timing fold change profiles constructed for Sniper results (top) and Rao et al. (2014) subcompartments (bottom).

Importantly, we found that the Sniper outperforms the baseline Gaussian HMM (Fig. S2A) and the recently published Megabase (Fig. S2B) by using the original Hi-C subcompartment annotations in Rao et al. (2014) as a benchmark. Fig. 2C shows that Sniper significantly outperforms Megabase and the Gaussian HMM in all subcompartments. Most notably, Sniper accurately annotates B2 and B3 regions, whereas Megabase frequently confuses B2 and B3 (Fig. S2B).

We then compared the prediction from Sniper with histone mark ChIP-seq and DNA replication timing data in GM12878 (Fig. 2D) obtained from ENCODE (Consortium, 2012). We determined enrichment of different epigenetic marks in each Sniper subcompartment and Rao et al. (2014) subcompartment by following the procedure in the Supplement Section V.b.2 from Rao et al. (2014). Overall we found that the enrichments with histone marks and replication timing are very consistent with the results in Rao et al. (2014) (Fig. 2D). This further suggests the overall high concordance between the predictions from Sniper, which only uses downsampled data (10% of the original read pairs), and the original Hi-C subcompartment annotations from Rao et al. (2014).

We then determined the minimum Hi-C coverage level at which Sniper remains accurate. We also compared the accuracy of Sniper and the Gaussian HMM at various coverage levels to determine the point at which Sniper no longer outperforms the HMM. Sniper accurately predicts subcompartment annotations for cell types with at least 250 million Hi-C read pairs. At 200 million read pairs, we also found that Sniper outperforms the Gaussian HMM baseline, but remains far less accurate compared to the results based on 250 million read pairs (see Table S2).

### Sniper annotations in other cell types are supported by genomic and epigenomic data

Because subcompartment annotations from high-coverage Hi-C data are only available in GM12878, we cannot directly compare the Sniper predicted subcompartments in other cell types to the results based on high-coverage Hi-C. We therefore used functional genomic data in K562 and IMR90, where a large number of epigenomic datasets are available, to evaluate Sniper predictions. Fig. 3A is an example showing that Sniper recovered missing contacts from the the sparse low coverage inter-chromosomal contact map of IMR90, revealing much clearer compartmentalized contact patterns whose boundaries strongly correlate with shifts in functional genomic data. A1 and A2 regions generally have early replication timing and dense H3K27ac and RNA-seq signals, whereas regions in B1, B2, and B3 replicate later and have lower transcriptional activities (see Fig. S3 and Fig. S4).

**Figure 3:**
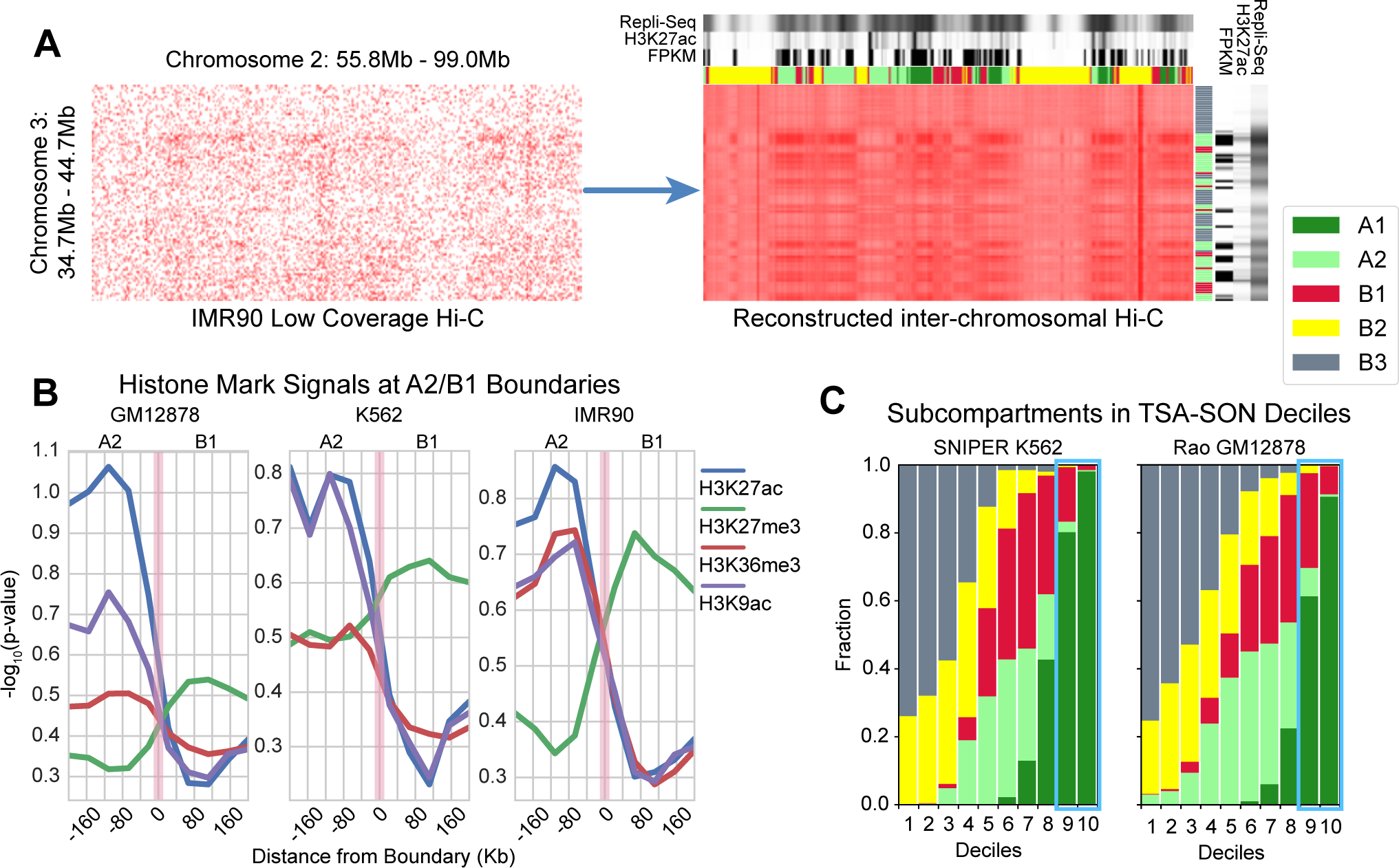
Sniper predictions correlate with various functional genomic data. **(A)** Reconstruction of the inter-chromosomal Hi-C contact matrix in IMR90. This example between chromosomes 2 and 3 shows that Sniper imputes missing contacts in the sparse matrix, recovers subcompartment-specific contact patterns, and predicts annotations that correlate with DNA replication timing Repli-seq, H3K27ac ChIP-seq, and RNA-seq (FPKM). **(B)** Relative histone mark signal p-value changes at the boundary between A2 (left) and B1 (right) in GM12878, IMR90, and K562. **(C)** Subcompartment distribution in K562 SON TSA-seq deciles for Sniper’s K562 subcompartments (left) and Rao et al. (2014)’s GM12878 subcompartments (right).

In addition, we observed significant shifts of histone mark signals in 400kb neighborhoods around subcompartment boundaries between A2 and B1 (Fig 3B for results in GM12878, K562, and IMR90) and other subcompartment boundaries (see Fig. S5). We focus on A2 to B1 transitions as they are usually the most frequent amongst the cell types in our analysis, e.g., about six times more frequent than A1 to B1 transitions. Furthermore, A2 and B1 are associated with euchromatin and facultative heterochromatin, respectively (Rao et al., 2014), and can both be transcriptionally relatively active. As a result, the large shift in histone mark signals across their boundaries signifies Sniper’s ability to differentiate spatially close and functionally similar subcompartments. Active marks such as H3K9ac, H3K27ac, and H3K36me3 are generally more enriched in A2 than B1 with a dramatic drop moving across the boundary, consistent with the significantly lower enrichment of active marks in B1 (see Fig. S2C), whereas the facultative heterochromatin mark H3K27me3 becomes more enriched across the A2-B1 boundary. These patterns of changes in epigenomic signals at the boundaries of subcompartments are consistent with changes in histone mark signals at subcompartment boundaries shown by Rao et al. (2014) and more recently by Chen et al. (2018), and the average log ratio between two epigenomic signals shown by Robson et al. (2017). We also observed changes of histone mark signals around A2 and B1 boundaries in downsampled GM12878, K562, and IMR90 using subcompartment annotations from Gaussian HMM clustering (Fig. S6). Compared to what we observed from the predictions based on Sniper, the patterns of signal changes around A2 and B1 boundaries in GM12878 and IMR90 based on Gaussian HMM are similar. However, signals in K562 annotated by Gaussian HMM showed no noticeable difference around A2 and B1 boundaries. This observation suggests that Gaussian HMM may not be appropriate to identify subcompartments with accurate boundaries for all cell types.

We found that genomic regions replicate much earlier in A1 and A2 subcompartments than in B subcompartments (Fig. S7A) in GM12878, K562, and IMR90. In addition, it is known that the level of histone modification of H3K27ac is associated with enhancer activities (Creyghton et al., 2010) and sometimes also transcriptionally active inter-LADs (van Steensel and Belmont, 2017). We found that H3K27ac generally has much higher signal in predicted A1 and A2 than in B compartment regions, and is virtually absent in predicted B2 and B3 (see Fig. S7B). Higher H3K27ac signals in B1-annotated regions suggest less transcriptional activity than regions in A1 and A2 but more activity than B2 and B3. Intermediate levels of transcriptional activity and increased abundance of H3K27me3 in the predicted B1 regions are indicative of its association with facultative heterochromatin (van Steensel and Belmont, 2017) (see Fig. S7C).

The recently developed TSA-Seq technique can reveal cytological distance between chromosomal regions to specific subnuclear structures (Chen et al., 2018). We compared Sniper subcompartments with TSA-seq scores (Fig. 3C). We used SON and LaminB TSA-seq that measures the distance to nuclear speckles and nuclear lamina in K562, respectively (note that TSA-seq is not available for other cell types studied in this work). We found that SON TSA-Seq signal shows significant stratification of the predicted subcompartments in K562. In particular, the highest TSA-SON decile is almost exclusively associated with A1 regions in the Sniper annotations. In contrast, using Rao et al. (2014) GM12878 annotations, a significantly higher portion of the highest decile is associated with B1 regions. Furthermore, none of Sniper’s predicted A2 regions are binned into the lowest 2 deciles while Rao et al. (2014)’s A2 regions are present in all low deciles. A scatter plot of SON TSA-seq and LaminB TSA-seq percentiles (Fig. S8) shows that almost all subcompartments tend to cluster better based on Sniper K562 subcompartment annotations instead of the original GM12878 subcompartment annotations. We found that in general Sniper annotations in K562 are partitioned into subcompartments with narrower TSA-SON and TSA-LaminB signal ranges compared to Rao et al. (2014) annotations in GM12878. These results suggest that the Sniper subcompartment annotations in K562 are accurate and offer a more appropriate comparison with SON and LaminB TSA-seq in K562 than the Rao et al. (2014) GM12878 subcompartment annotation (which was the approach Chen et al. (2018) used).

### Sniper facilitates the identification of subcompartment patterns across different cell types

We next applied Sniper to predict Hi-C subcompartments in K562, IMR90, HeLa, HUVEC, HMEC, HSPC, T Cells, and HAP1. Together with the subcompartments in GM12878, this allows us to perform a detailed comparison of subcompartment conservation and changes across multiple cell types. 100kb genomic regions are partitioned into 13 conservation states (see Methods for detailed definitions) based on each region’s subcompartment annotation distribution among nine cell types. States are termed states 1-12 and NC, sorted by ascending entropy of cross cell type annotations, of which state 1 has the lowest entropy and refers to genomic regions with the most conserved cross cell type annotations, states 2-12 gradually increase in entropy and decrease in conservation, and state NC refers to the dynamic non-conserved state. Conservation states 2, 5, 7, and 8 also denote genomic regions that contain cell type specific subcompartment annotations. States 1-3 occupy large fractions of the genome, indicating that a large portion of the genome contains relatively conserved subcompartment annotations (Fig. 4A). Notably, the A1 subcompartment appears to be the most conserved subcompartment across cell types, with about 40% representation in the most conserved state 1. By contrast, there is less B1 presence in the more conserved states, consistent with the observations in Rao et al. (2014) that B1 is associated with facultative heterochromatin. The NC state comprised about 15% of the genome and contained relatively few A1 and B3 regions, suggesting A1 and B3 may be more conserved across cell types than other subcompartments.

**Figure 4:**
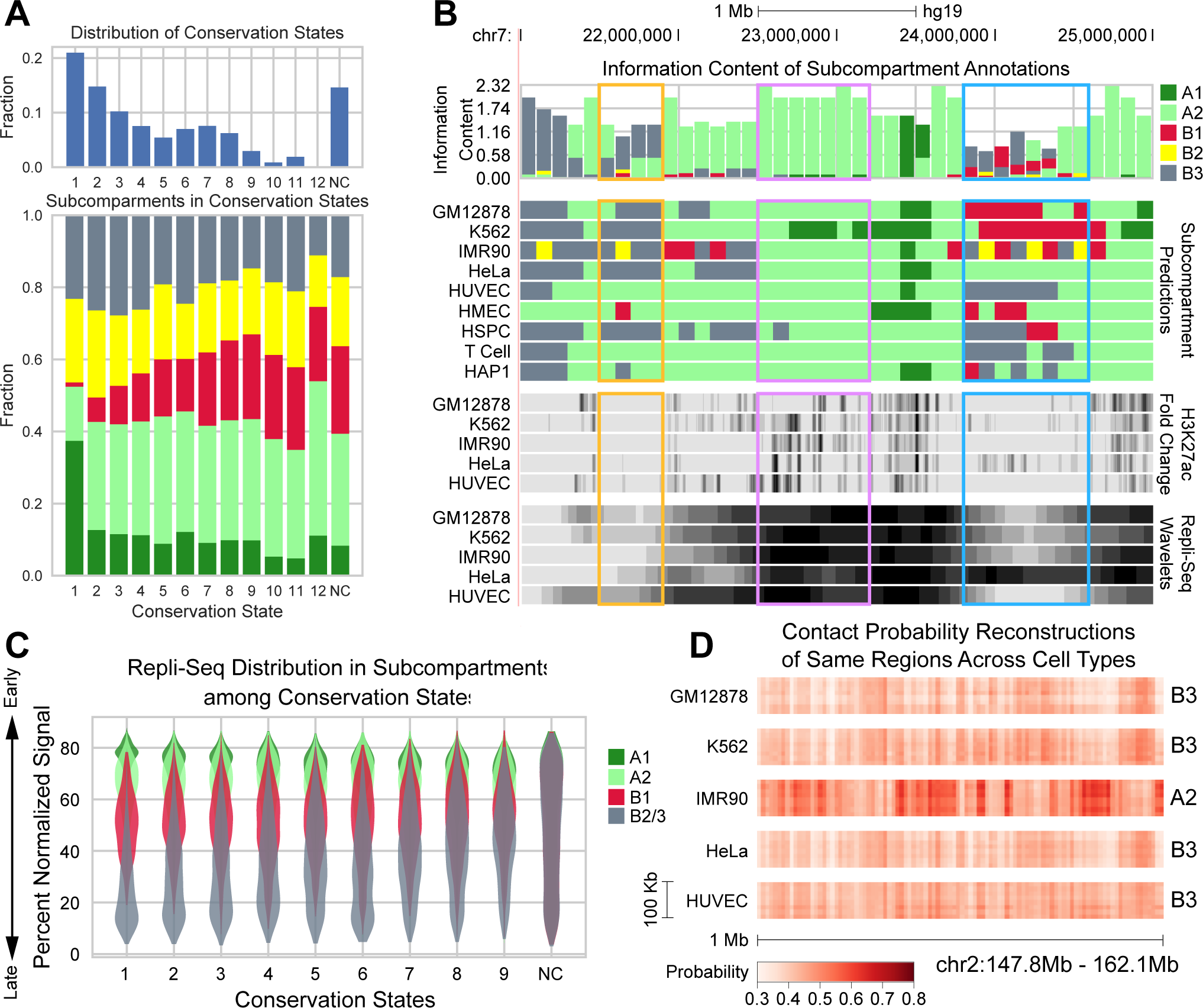
Sniper allows comparisons of subcompartments across different cell types. **(A)** (Top) Distribution of thirteen conservation states in the genome. (Bottom) Distribution of subcompartment regions in each conservation state. **(B)** UCSC genome browser shot displaying the information content (IC), cross cell type predictions, H3K27ac fold change, and smoothed Repli-seq wavelets of each 100kb region. Subcompartments offering the most information, associated with taller bars in the IC track, are more conserved across nine cell types. **(C)** For each conservation state, we show Repli-seq signal distribution in the most frequent subcompartment annotations among nine cell types at each 100kb region. Regions in conservation state 1 are the most conserved, reflected by low variance of Repli-seq signal in each subcompartment. Regions in the NC state are the most dynamic, suggesting multiple annotations for a single chromatin region and high variance in Repli-seq signal. **(D)** Hi-C reconstructions across cell types in chr18 (47.1-47.9Mb) where IMR90 is A2-specific and other cell lines are predicted as B3.

Information content (see Methods), H3K27ac ChIP-seq fold change, and smoothed Repli-seq signals at each region of the Genome Browser shot (Fig. 4B) show conserved and dynamic functional genomic patterns across cell types. Similar to information content of position weight matrices for transcription factor binding motifs, subcompartment information content reflects the information gained from annotations across all cell types in 100kb genomic bins. Genomic regions with high information content have significantly more conserved annotations across cell types than regions with low information content. Conserved A2 regions, shown in the purple segment of Fig 4B, can be expected to retain conserved annotations and functional genomic patterns even in cell lines not in our analysis. Less informative regions (Fig 4B, yellow and blue) exhibit inconsistent functional genomic signals. Regions with HeLa-specific A2 annotations (Fig 4B, blue rectangle) show increased abundance of H3K27ac signal and much earlier replication timing compared to other cell lines. These regions are annotated as B1, B2, and B3 in other cell types and correspond to lower H3K27ac signals and later replication timing.

The amount of information gained in each conservation state is reflected in the Repli-seq distribution in subcompartment modes across states (Fig. 4C). For each region in a conservation state, its cross cell type Repli-seq signals were binned according to the region’s mode, defined as the most frequent subcompartment annotation among 9 cell types. We then plotted the violin plots of Repli-seq signals in each mode of the conservation state. We binned Repli-seq signals for all other conservation states except states 10, 11, and 12, which contained too few 100kb regions. We found that more conserved states show less variance of Repli-seq signals in each mode because cross cell type predictions are less varied. Less conserved states such as states 8 and 9 exhibit much higher Repli-seq variance in each mode, especially B1 and B3. Repli-seq distributions of all modes virtually overlap in the NC state, further showing high variance of functional genomic signals in more dynamic subcompartment regions across cell types. Because Repli-seq is virtually identically distributed in B2 and B3, the two subcompartments are merged in Fig. 4C.

Hi-C reconstructions at genomic regions with cell type specific annotations are distinct from the same regions in other cell types. Fig. 4D shows an example of A2 regions specific to IMR90 that exhibit significantly more frequent contacts compared to the same region in other cell types, which are annotated as B3.

Taken together, these results demonstrate that Sniper provides us with the capability to reliably compare Hi-C subcompartment annotations in multiple cell types and analyze cross cell type patterns of conservation and variation of Hi-C subcompartments.

### Conserved and cell type specific subcompartment patterns show distinct gene functions

Genomic regions where transcription activity is high in a single cell type may reveal genes that contribute to unique cellular functions. Here we focus on cell type specific A2 regions instead of A1 even though both subcompartments have high transcription activity. Cell type specific A1 regions are frequently annotated as A2 in other cell types, and therefore likely to show high transcription activity across multiple cell types, obscuring cell type specific gene activity. Conversely, cell type specific A2 regions are much less frequently annotated as A1 in other cell types (see Fig. S9), and are more likely to reveal cell type specific gene activity. Such regions are much less likely to share high transcription activity across multiple cell types. Gene Ontology (GO) biological processes associated with constitutive genes (Table. 1) may reveal housekeeping processes required for basic cellular functions. The most enriched biological processes from GO enrichment analysis include catabolism, translation, protein targeting, molecular transport, and RNA metabolism among others. Furthermore, log_10_ *p*-values of enriched processes show that genes in constitutive A2 regions contain significant housekeeping functions.

**Table 1:** GO term enrichment of genes in genomic regions with conserved A2 annotations across cell types and cell type specific A2 annotations in T cells and K562.

| **Biological Process (Constitutive A2)** | **log<sub>10</sub> p-value** |
| --- | --- |
| SRP-dependent cotranslational protein targeting to membrane | -27.6498 |
| mRNA metabolic process | -26.1152 |
| translational initiation | -25.6716 |
| nuclear-transcribed mRNA catabolic process | -22.2874 |
| cellular localization | -21.5243 |
| peptide transport | -17.4056 |
| cellular amide metabolic process | -16.1778 |
| macromolecule localization | -16.163 |
| nitrogen compound transport | -15.5272 |
| cellular protein metabolic process | -15.1904 |
| **Biological Processes (T Cell Specific A2)** |  |
| Immune system process | -8.523 |
| T cell activation | -6.319 |
| **Biological Processes (K562 Specific A2)** |  |
| Ribonucleoprotein complex biogenesis | -5.355 |
| rRNA metabolic process | -3.3224 |

We then compiled cell type specific A2 annotations across 9 cell types and highly expressed genes in cell type specific A2 regions. We divided genes into cell type specific A2 sets (see Methods), used the method in (Reimand et al., 2007) to obtain GO terms associated with genes in each set, and removed redundant GO terms using REVIGO (Supek et al., 2011). Cell type specific subcompartment annotations can provide a clear picture of functions and pathways enriched in certain cell types such as T cells. Specifically, genes in T cell-specific A2 regions are highly associated with the immune system process and T cell activation, functions characteristic of T cells. However, genes in facultative A2 regions in other cell types do not necessarily produce enrichment that explains cell type specific functions. RNP complex biogenesis and rRNA metabolic process are among the most highly enriched processes in K562, but are also significantly enriched in constitutive A2. However, a single cell type specific region may contain many genes that act as passenger genes to a small number of key genes for driving such spatial localization changes of chromatin in nucleus (e.g., Khanna et al. (2014)), obscuring cell type specific functions and pathways.

## Discussion

In this work, we introduced Sniper, a new computational method that imputes inter-chromosomal contacts missing from sparse Hi-C datasets and predicts subcompartment annotations at 100kb scale across multiple cell types. We found that Sniper annotated subcompartments in the GM12878 with high accuracy and outperformed a state-of-the-art method, Megabase. In GM12878, K562, IMR90, HeLa, HUVEC, HMEC, HSPC, T cells, and HAP1, we showed that Sniper predictions correlate well with functional genomic data including histone marks, replication timing, RNA-seq, and TSA-seq. Genomic regions with conserved Sniper annotations across these 9 cell types occupy a significant portion of the genome and shared similar abundance of epigenomic signals. Regions with constitutive A2 predictions are generally associated with housekeeping functions and pathways. Cell type specific A2 predictions correlate with biological processes specific to some cell types.

Sniper is able to achieve accurate subcompartment annotations in cell types with Hi-C coverage as low as 250 million reads. In this study, Hi-C data for different cell types typically have more than 250 million Hi-C read pairs, between 400 million to 1 billion, suggesting that the Sniper subcompartment predictions are accurate. Compared to GM12878 annotations in Rao et al. (2014), we only need approximately 20 times fewer Hi-C reads to reliably annotate subcompartments using Sniper. Therefore, Sniper has the potential to significantly reduce the cost of Hi-C experiments to analyze subcompartments across many different cellular conditions.

The Hi-C subcompartment predictions from Sniper can be compared to results based on other analysis approaches and datasets. For example, we expect that the Sniper predictions of Hi-C subcompartments can be used to further validate and compare with results from polymer simulations (Sanborn et al., 2015; Nuebler et al., 2018), 3D genome structure population modeling (Tjong et al., 2016; Hua et al., 2018), and regulatory communities mining based on whole-genome chromatin interactomes (Dai et al., 2016). In addition, recently published new genome-wide mapping methods (Chen et al., 2018; Quinodoz et al., 2018; Beagrie et al., 2017) may provide additional training data other than Hi-C, as well as experimental data validation to improve our method.

Currently Sniper is limited by its training data, which contains about 500 million mapped read pairs from the original Hi-C reads in GM12878. The Hi-C coverage of other cell lines tends to vary, which can impact Sniper’s overall accuracy when applied to some cell lines. As a result, Sniper could incorrectly annotate some regions in a cell line if Hi-C coverage is too high or too low. In addition, the ratio between intra-chromosomal and inter-chromosomal reads can vary across cell lines, which we did not explicitly control for. This ratio could exhibit high variance across different cell types and influence the accuracy of Sniper’s predictions. Future work should make Sniper more coverage invariant and produce consistent annotations regardless of the Hi-C coverage of its inputs. In addition, the Hi-C subcompartment annotations used in Sniper are also largely relying on the original annotations in GM12878 from Rao et al. (2014). Although the results in this work demonstrate that these subcompartment definitions may well represent primary subcompartments in many cell types, it is also possible that some cell types may have their distinct subcompartment organizations. Future work can be performed to train Sniper to categorize genomic regions into different sets of subcompartments not limited to the five primary subcompartments used in this work. Furthermore, the inability to isolate cell type specific functions in some cell types suggests more work should be done to determine genes in facultative subcompartment regions that most significantly contribute to cell type specific spatial localization and function. Nevertheless, this work demonstrated that Sniper has the potential to become a useful tool to offer new perspectives of 3D genome organization changes in different cell types.

## Methods

### The denoising autoencoder for inferring high resolution inter-chromosomal Hi-C contacts

We aim to recover missing contacts from sparse inter-chromosomal Hi-C contact maps by constructing a denoising autoencoder (Vincent et al., 2008) (Fig. 1B), which uses rows of the downsampled Hi-C contact probability matrix (see later section) in GM12878 as inputs, and targets the corresponding rows of the dense matrix. Each row of the matrix is a vector of contact probabilities between one locus in an odd (or even) chromosome and all loci in the even (or odd) chromosomes. The denoising autoencoder in Sniper contains a total of 9 sequential layers with *N*_*loci*_, 1024, 512, 256, 128, 256, 512, 1024, and *N*_*loci*_ neurons, respectively, where *N*_*loci*_ refers to the number of rows or columns in the input contact matrix. Layers with *N*_*loci*_ neurons are the input and output layers, the layer with 128 neurons is the latent layer, and the remaining layers are hidden layers. The autoencoder network is trained using a moderate coverage Hi-C matrix obtained by randomly removing 90% of the original GM12878 Hi-C read pairs to reflect the sparsity and coverage levels of other cell types. We input a subset of *N* total rows into the autoencoder’s input layer. Its output layer targets corresponding rows in the high coverage Hi-C matrix. The layers of the encoder and decoder, pertaining respectively to the layers before and after the autoencoder’s latent layer, contain 12-14 million parameters, approximately 10% of the *N* × *N*_*loci*_ ≈ 145 million sparse inputs in the training matrix. The 128 dimension of the latent layer limits the number of parameters in the autoencoder to approximately match our downsampling ratio of 1:10, and enables the downstream classifier to accurately predict subcompartment annotations.

#### Linear transformations to compute neural layer outputs

We developed a denoising autoencoder that is comprised of linear layers with neurons whose activations are computed by:

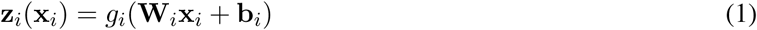

where **z**_*i*_ is the activated output of layer *i*, **x**_*i*_ is the *n*-dimensional input to layer *i*, **W**_*i*_ is the *m* × *n*-dimensional weight matrix (where *m* is the layer’s output dimensionality) of layer *i*, **b**_*i*_ is the *m*-dimensional bias vector of layer *i*, and *g*_*i*_ is the activation function applied element-wise to the output vector for layer *i*.

#### Nonlinear activation to promote separability and consistency of autoencoder outputs

We apply rectified linear unit (ReLU) activation (Nair and Hinton, 2010) to the hidden layers:

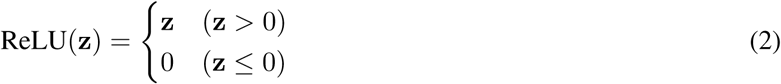

where all non-positive values in the output vector **z** are set to 0 to introduce sparsity. Sparse neural activation is less entangled, more linearly separable, and more efficiently propagates information throughout the network. In addition, ReLU has been shown to be suitable for naturally sparse data (Glorot et al., 2011).

Of the hidden layers, those with 1024 and 256 neurons are forwarded into 25% dropout layers to reduce overfitting (Witten et al., 2016). The latent and output layers are activated linearly and sigmoidally, respectively, with no dropout:

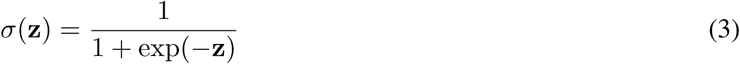

where the values in the activated output *σ*(**z**) are constrained between 0 and 1 as the exponential converges to 0 and 1 (as values in a layer’s output *z* go to −∞ and +∞). The latent layer is linearly activated to maximize the encoding space that latent variables can occupy. The output layer is sigmoidally activated to match the range of values in the input probability matrix.

#### Binary cross-entropy to optimize the autoencoder

We use binary cross-entropy (BCE) loss to assign weights to samples whose autoencoder output values deviate significantly from corresponding target values:

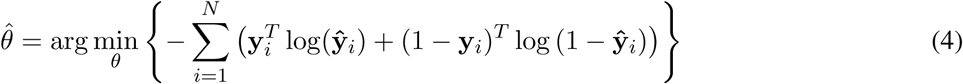

where the autoencoder parameters *θ* are optimized to minimize the cross entropies between model outputs **ŷ**_*i*_ and target outputs **y**_*i*_ for all training inputs *i* ∈ {1, …, *N*}. The autoencoder can also be optimized using mean-squared error loss with little difference in performance (Creswell et al., 2017).

For implementation, gradients of the weights and biases in the model are computed using backprogation (Rojas, 1996) and weights are updated using the RMSProp (Hinton et al., 2012). The autoencoder is trained for 25 epochs using a batch size of 32 and learning rate of 0.001.

The training set contains the first 7,000 rows in the contact probability map, which includes genomic loci in chromosomes 1, 3, 5, and 7, occupying about 55% of all loci in odd-numbered chromosomes. The remaining rows in the contact probability map form the test set. We found that using different sets of chromosomes did not significantly affect the recovery of high coverage Hi-C data and the annotation of subcompartments (see Fig. S1). In addition, the autoencoder was re-trained using even-numbered chromosomal regions as training inputs. This new training set includes loci in chromosomes 2, 4, 6, 8, and 10 and occupied about 60% of loci in even-numbered chromosomes. We transpose the downsampled Hi-C matrix, compute its probability maps, and follow the same training process, targeting the transposed high coverage Hi-C probability map. The re-trained autoencoder model outputs the same Hi-C map as the initial model (see Fig. S10). The size of the input and output layers is adjusted to equal the number of contacts between each even-numbered chromosomal region and all odd-numbered regions.

### The classifier for predicting Hi-C subcompartment annotations

We developed a multi-layer perceptron model with two hidden layers to classify latent representations of inter-chromosomal contacts into subcompartment assignments (Fig. 1C). The MLP network contains layers with 128, 64, 16, and 5 neurons. The 128-neuron layer pertains to the input layer, the 64- and 32-neuron layers are the hidden layers, and a 5-neuron layer is the output layer (corresponding to five primary subcompartments). The network is trained using the latent representations of inter-chromosomal contacts and the corresponding GM12878 subcompartment labels from Rao et al. (2014). We then input the 128-dimensional representations of genome-wide contacts in other cell types into the trained classifier to infer their subcompartment annotations.

Sigmoid activation is applied to the input latent variables, limiting input values between 0 and 1 and mitigating bias towards high numerical inputs. We apply ReLU activation to the output of each hidden layer, which will subsequently be forwarded to 25% dropout layers. The output layer contains 5 neurons (each representing a possible subcompartment annotation) which are activated with softmax to ensure that subcompartment probabilities summed to 1:

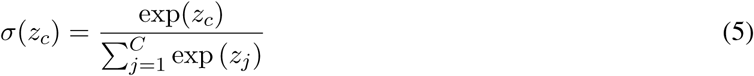

where the exponential activation of a class *c*, exp(*z*_*c*_), is normalized by the sum of exponential activation across all classes 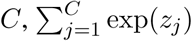. The output likelihoods indicate the most likely annotation of a 100kb genomic bin.

The training set is balanced (see Methods below) to ensure that each subcompartment is equally represented in the training set. Our classifier uses a balanced set of latent representations of the same loci used to train the autoencoder as inputs and targets their corresponding subcompartment annotations **y** based on high coverage Hi-C in Rao et al. (2014). We validated the model by comparing the predicted annotations of the remaining loci’s latent variables to the Rao et al. (2014) annotations. The model is optimized using categorical cross-entropy loss between predicted and target outputs:

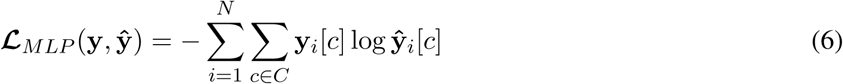

where ŷ is the predicted output, **y** is the target output, and *c* ∈ *C* are the possible output classes. The loss function sums over the class-specific entropy loss **y_i_**[*c*] log **ŷ**_*i*_ for all classes in each training sample *i* ∈ {1, …, *N*}. The weights in the classifier are updated by computing gradients of the loss function with respect to the weights:

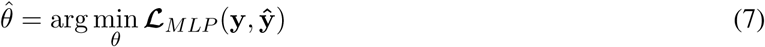

where *θ* is the set of model weights. Each epoch’s learning rate is adjusted using RMSProp (Hinton et al., 2012). Two independent classifiers are trained to annotate regions in odd- and even-numbered chromosomes.

### Converting Hi-C contact maps into Hi-C contact probabilities

We converted Hi-C contacts into contact probabilities to mitigate the effects of extreme Hi-C signals and enable neural networks to use binary cross-entropy loss. Eq. 8 was applied element-wise to an inter-chromosomal Hi-C map, returning a matrix of contact probabilities *P*_*ij*_ constrained between 0 and 1.

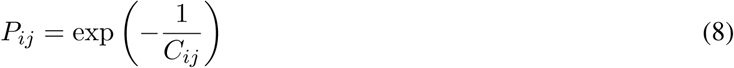

where *C*_*ij*_ refers to the contact frequency between genomic loci *i* and *j*. Contacts probabilities with values constrained between 0 and 1 allow a neural network’s weights to be optimized using BCE loss instead of mean-squared-error (MSE) loss and mitigate the effects of extreme outliers. High frequency chromatin contacts can disproportionately influence activation in a neural network with linear neurons, leading to incorrect chromatin subcompartment annotations and a less robust neural network. Even after log-normalization in Rao et al. (2014), Sniper can still become skewed by the logarithms of extreme values. Furthermore, the value range of the input exceeds the (0, 1) range and pushes Sniper to compute gradients derived from MSE loss. MSE loss optimizes for mean-squared distance between network outputs and targets, which can result in regression to the mean of the targets. Using BCE loss retains Hi-C contact patterns in the autoencoder output. Sniper’s goal is to capture contact frequency patterns in order to infer subcompartment annotations, making optimization for patterns far more important than optimizing for mean-squared distance. In addition, extreme outliers in the contact matrix will have corresponding contact probabilities that converge to 0 or 1, values which will introduce much less bias into the autoencoder. While Sniper’s inputs could also be constrained between 0 and 1 by applying a sigmoid function to the input layer activation, the input would have to be further balanced by training an additional set of weights and biases.

Probability maps were computed for cell types GM12878, K562, IMR90, HeLa, HUVEC, HMEC, HSPC, T-Cells, and HAP1. Because GM12878’s Hi-C coverage is much higher compared to the other cell types, we simulated the sparsity of other cell types’ Hi-C maps by downsampling GM12878’s inter-chromosomal matrix by 1:10 and computing its downsampled probability map. The downsampled probability matrix serves as the training input for the Sniper autoencoder and GM12878’s dense matrix serves as its target output during training.

### Training set balancing

The training set of the classifier was balanced so that each subcompartment was equally represented to remove bias towards specific subcompartments. We set the number of samples per subcompartment to be a number *N* that is greater than the number of regions in the most common subcompartment in the GM12878 training set. We then define an array *B* corresponding to the balanced training set containing 5 × *N* training samples – *N* samples per subcompartment.

For each of the five primary subcompartments *c*, we randomly sample two latent variables *x* and *y* of chromatin regions that belong to subcompartment *c*. We subsequently compute *r*, a vector novel to the training set whose values lie at a random point in between the values *x* and *y*, i.e.,

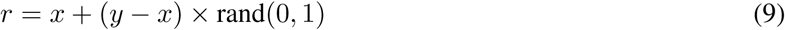

where rand(0, 1) is a random variable sampled from a uniform distribution between 0 and 1. We then append *r* to *B* and repeat random sampling for *N* − 1 iterations. *N* random samples are then taken for each of the remaining subcompartments.

### Methods of comparing Sniper results in different cell types

#### Transition of histone marks near subcompartment boundaries

Epigenomic marks can serve as indicators of the overall accuracy of predicted annotations, even though they are not perfectly predictive of subcompartment state. We compiled histone marks ChIP-seq *p*-values in genomic regions within 400kb of subcompartment boundaries, defined as nucleotide positions where subcompartment annotations of adjacent 100kb chromatin regions are different.

#### Conserved and dyanmic subcompartment annotations across multiple cell types

In this work, Sniper is applied to 9 cell lines – GM12878, K562, IMR90, HeLa, HUVEC, HMEC, HSPC, T cells, and HAP1 – to determine regions whose subcompartment annotations are conserved and dynamic across multiple cell types. We divided subcompartment annotations in thirteen conservation states based on the entropy of each 100kb region cross cell type annotations as follows:

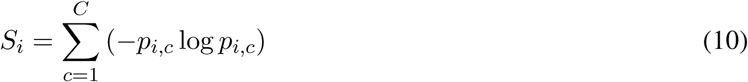

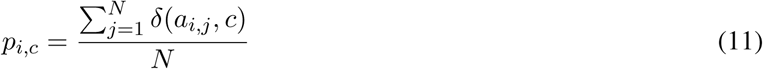

where *S*_*i*_ is the total entropy of region *i* subcompartment annotations, summed over the entropy of all *C* subcompartments. The fraction of subcompartment *c* at region *i*, *p*_*i,c*_, is computed by counting the number of occurrences of subcompartment *c* over all *N* cell types, 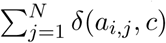, and dividing by the total number of cell types *N*. *δ* (*a*_*i,j*_*, c*) = 1 if the annotation *a*_*i,j*_ of cell type *j* is equal to *c* at region *i*.

Because annotations are discrete, Eq. 10 and 11 yielded 23 possible entropy values, each corresponding to a unique distribution of annotations across cell types. Of these 23 states, 11 were associated with fewer than 5 out of 9 cell types sharing the same subcompartment annotation. The 11 states without a majority subcompartment were merged into a single non-conserved (NC) state. We sorted the remaining 13 states in order of entropy, with the lowest entropy state 1 denoting the most conserved cross cell type regions, and the higher-numbered states denoting less conserved and more dynamic regions.

To represent subcompartment conservation and dynamics, we computed information content of each 100kb region. Information content was computed similar to entropy, but normalizing subcompartment-specific fractions by a background probability within the logarithm term:

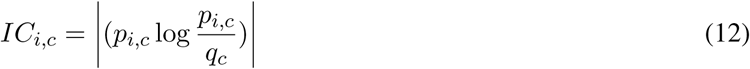

where *IC*_*i,c*_ is the information content of subcompartment *c* at region *i*, *p*_*i,c*_ is computed in Eq. 11, and *q*_*c*_ = 0.2 is the background probability of subcompartments assuming uniform subcompartment distribution. High information content corresponds to regions with more conserved annotations while low information content corresponds to more dynamic regions across cell types.

#### Isolating genes in regions with cell type specific annotations

Genomic regions in the A2 subcompartment have increased gene expression in general, but they are not guaranteed to have high gene expression, characterized by their overall but not universally high RNA-seq signals. Genes in regions with low RNA-seq signal are poorly expressed and can obscure the enrichment of Gene Ontology (GO) terms associated with highly expressed genes in cell type specific A2 regions. We instead used a combination of Sniper’s A2 annotations and FPKM from RNA-seq gene quantification to substantially decrease the size of gene sets in cell type specific A2 regions and reveal far more enriched GO terms. For each cell type, we sorted genes by their FPKM and added genes with FPKM above a threshold into our GO analysis set. The threshold depends on the number of cell type specific A2 regions in a cell type and gene density in these cell type specific regions, and was selected to limit the size of our gene sets to 2000.

### Hi-C data acquisition

The Hi-C data of GM12878, K562, IMR90, HeLa, HUVEC, and HMEC were obtained from Rao et al. (2014). The Hi-C data of HSPC, T Cell, and HAP1 was obtained from (Joeng et al., 2017). We used the Juicebox tool (Durand et al., 2016) to extract 100kb inter-chromosomal contacts from. hic files.

## Acknowledgement

K.X. is a predoctoral trainee supported by National Institutes of Health T32 training grant T32 EB009403 as part of the HHMI-NIBIB Interfaces Initiative. J.M. acknowledges support from the National Institutes of Health Common Fund 4D Nucleome Program grant U54DK107965, National Institutes of Health grant R01HG007352, and National Science Foundation grant 1717205. The authors would like to thank Andrew Belmont and members of Jian Ma’s laboratory (Yang Zhang, Ruochi Zhang, Yuchuan Wang, Ben Chidester, Yang Yang, Ashok Rajaraman, and Zhenghui Wang) for suggestions.

## Author Contributions

Conceptualization, J.M.; Methodology, K.X. and J.M.; Software, K.X.; Investigation, K.X. and J.M.; Writing – Original Draft, K.X. and J.M.; Writing – Review & Editing, K.X. and J.M.; Funding Acquisition, J.M.

## Declaration of Interests

The authors declare no competing interests.

