## Supplemental Information for "Revealing Hi-C subcompartments by imputing high-resolution inter-chromosomal chromatin interactions"

### Supplementary Figures and Tables

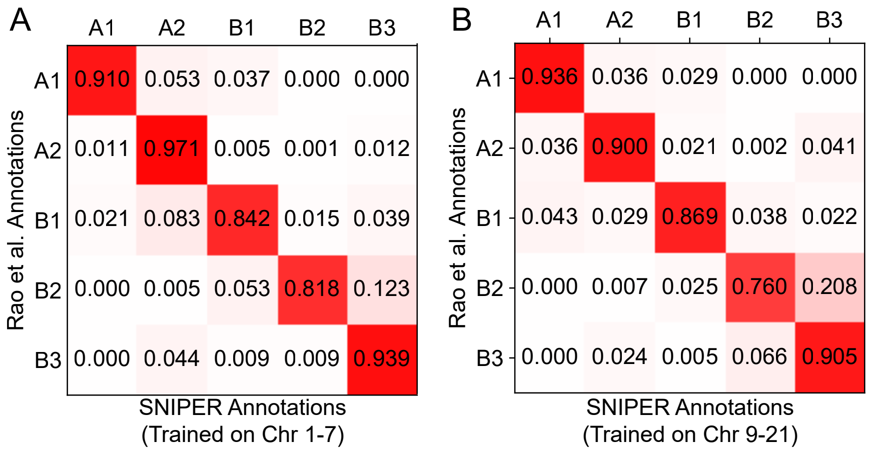


**Figure S1:** **A**. Accuracy of SNIPER when trained using rows in chromosomes 1, 3, 5, and 7. **B**. SNIPER's accuracy when trained on rows in chromosomes 9, 11, 13, 15, 17, 19, and 21.

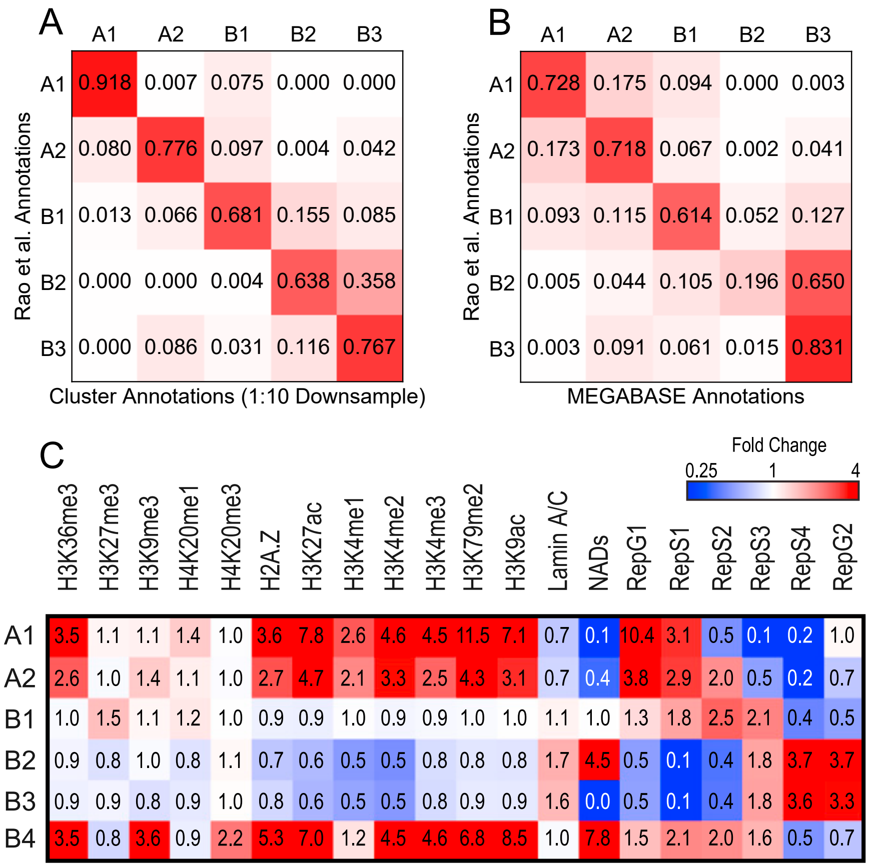


**Figure S2:** **(A)** Confusion matrix between Rao et al. (2014) subcompartment annotations and annotations from a Gaussian HMM clustered on downsampled GM12878 (490M reads) Hi-C matrix. **(B)** Confusion matrix between Rao et al. (2014) annotations and MEGABASE annotations. **(C)**.Epigenetic marker enrichment in each Rao et al. (2014) subcompartment.

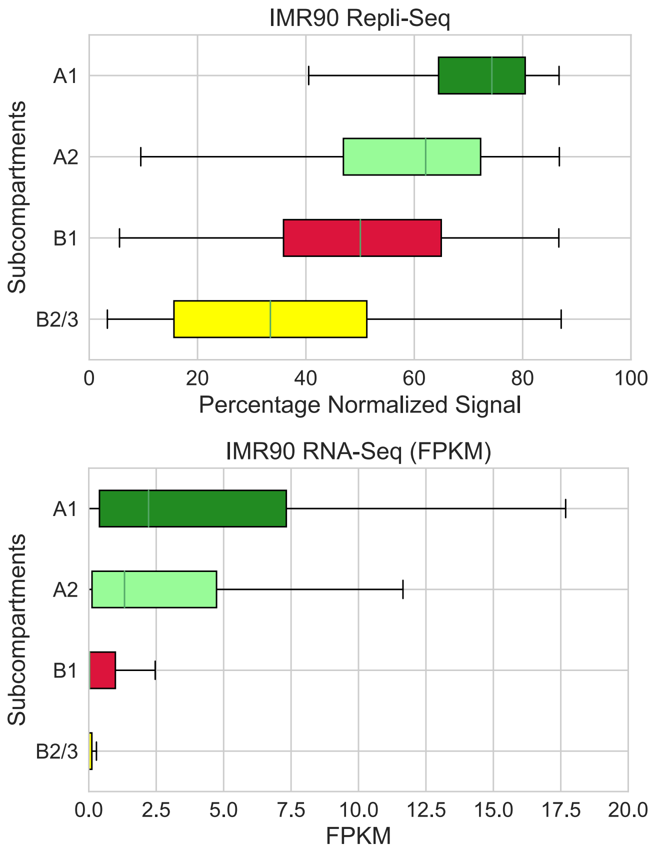


**Figure S3:** Distribution of Repli-seq and RNA-seq FPKM for each subcompartment in IMR90.

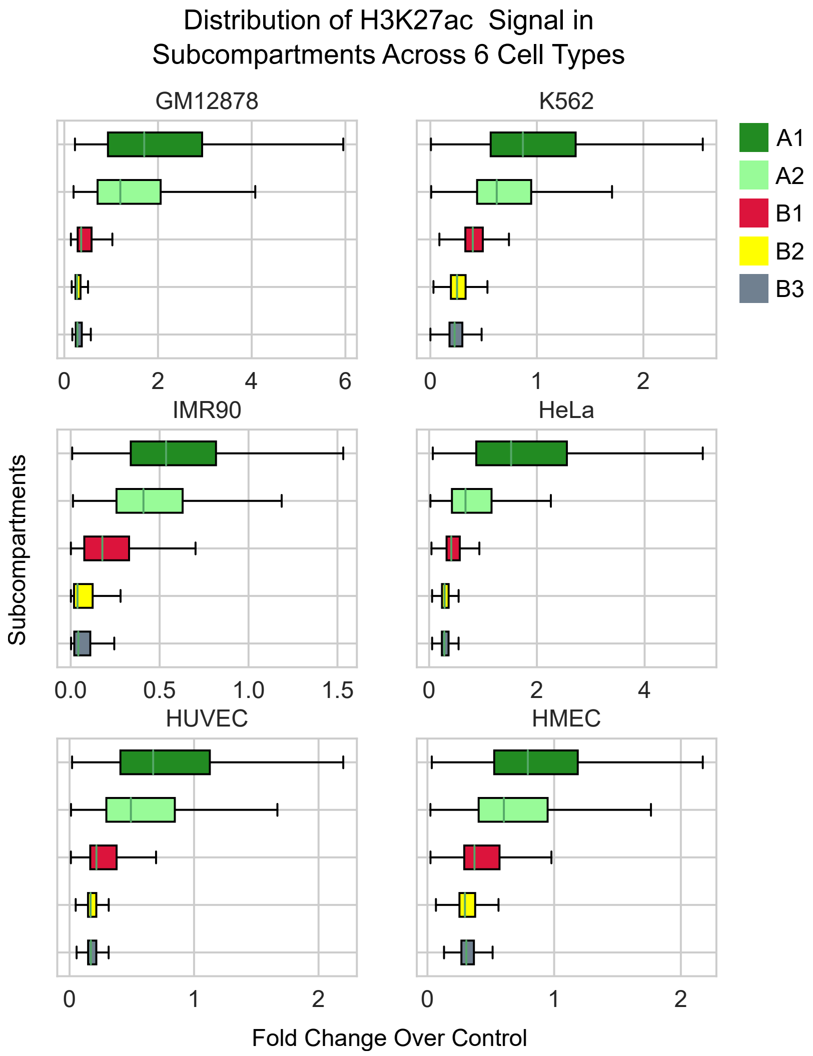


**Figure S4:** Distribution of H3K27ac fold change over control for each subcompartment in GM12878, K562, IMR90, HeLa, HUVEC, and HMEC.

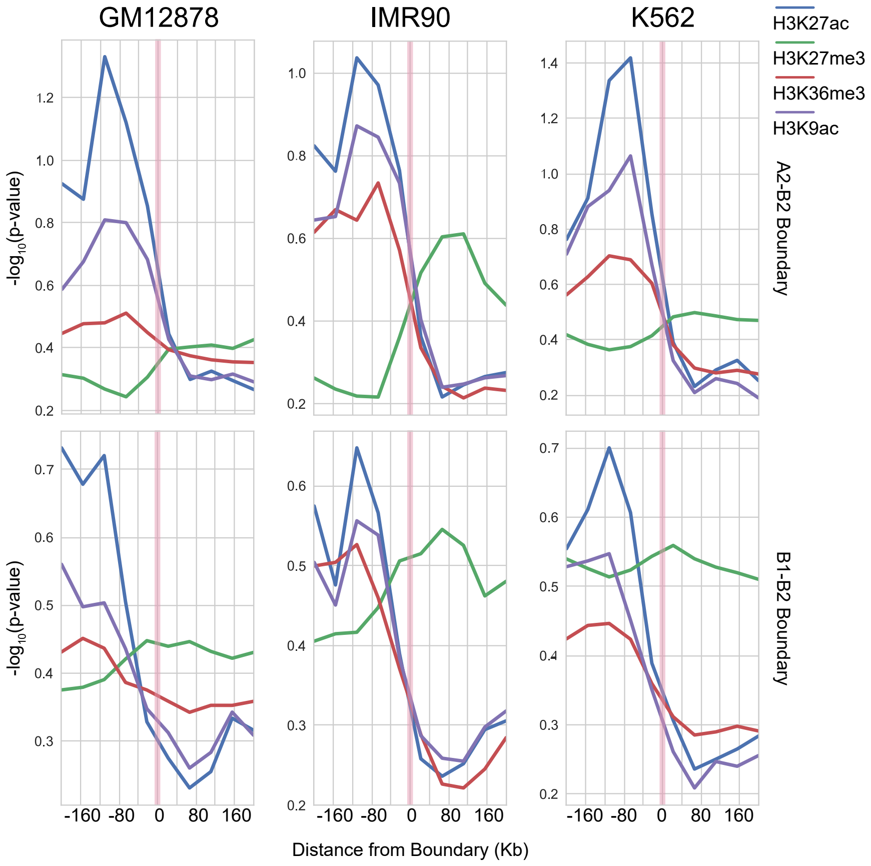


**Figure S5:** Histone mark signals across the A2-B2 and B1-B2 boundaries in GM12878, IMR90, and K562.

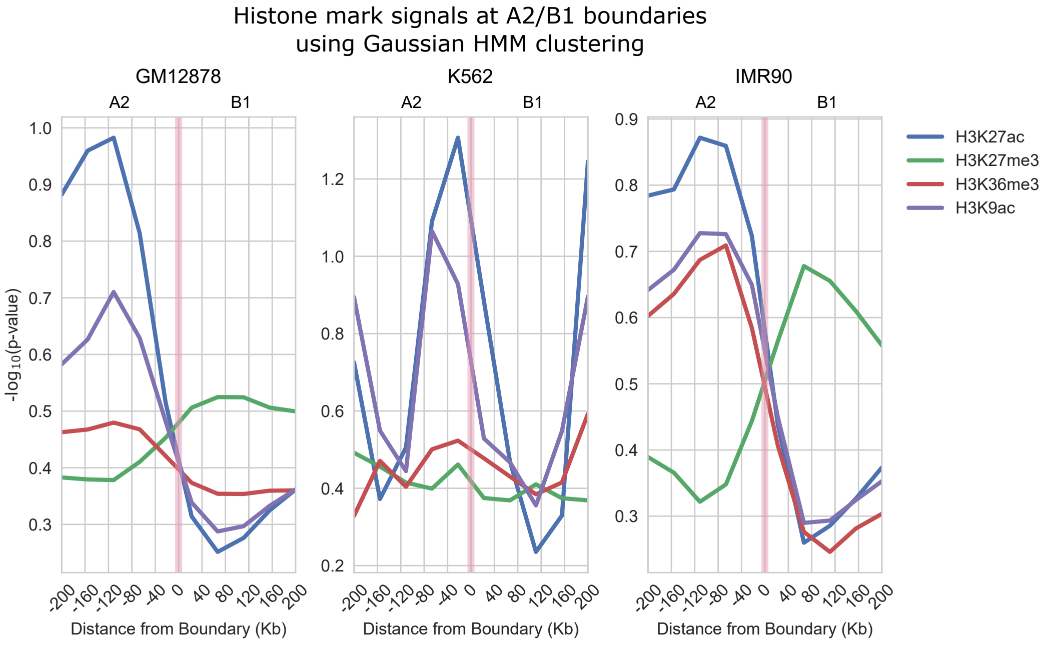


**Figure S6:** Histone mark p-values across the A2 and B1 boundary using subcompartment annotations obtained from Gaussian HMM clustering.

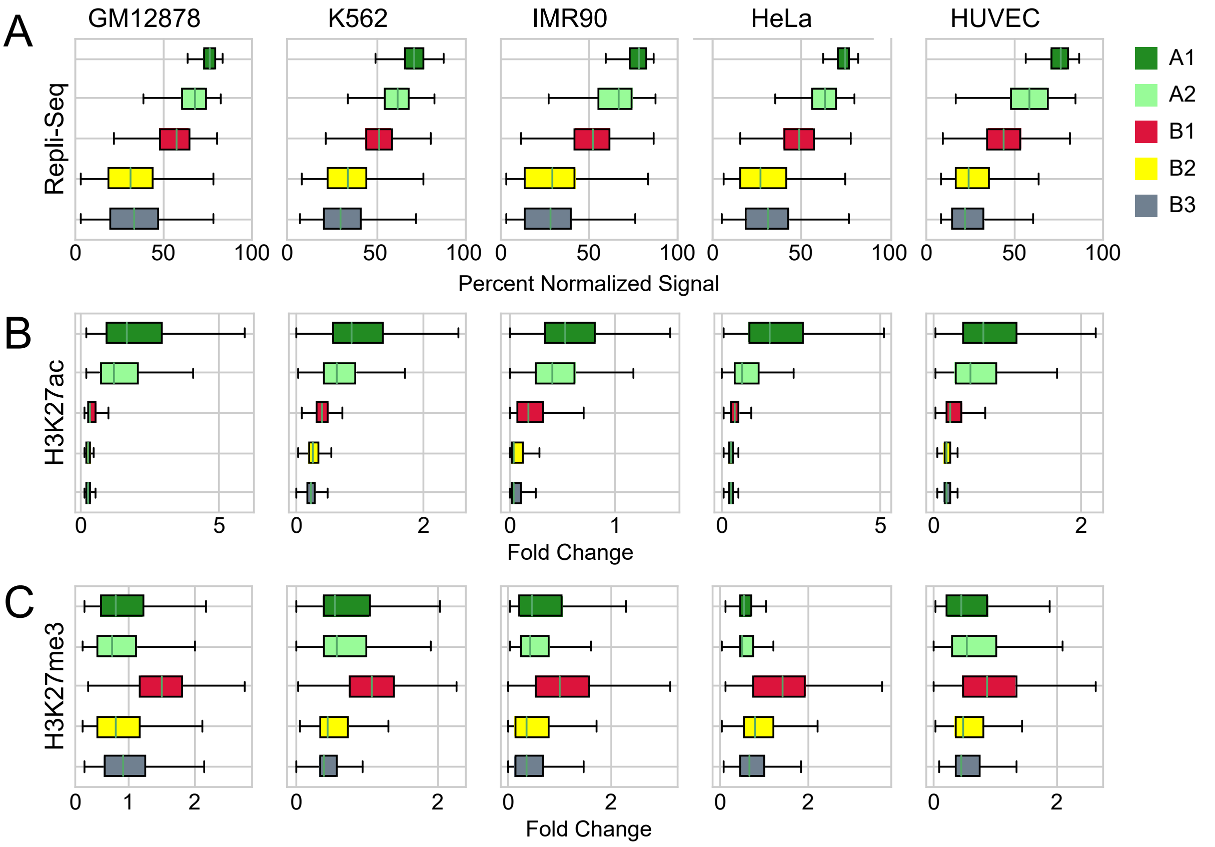


**Figure S7:** ChIP-seq fold change of H3K27ac and H3K27me3 and Repli-seq in predicted subcompartments across cell types GM12878, K562, IMR90, HeLa, and HUVEC.

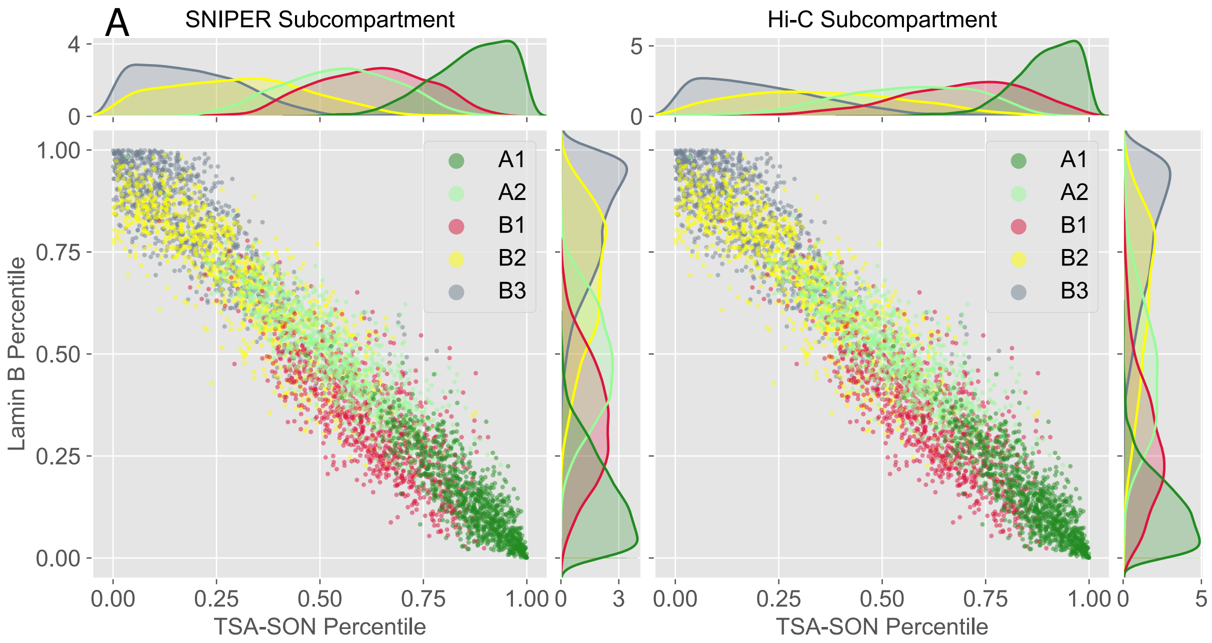


B

Variance in each subcompartment ($\times 10^{-2}$)

| | | A1 | A2 | B1 | B2 | B3 |
|---|---|---|---|---|---|---|
| **SON** | SNIPER | 0.921 | 1.932 | 1.840 | 2.871 | 1.871 |
| | Rao et al. | 0.634 | 3.072 | 2.601 | 4.007 | 2.515 |
| **Lamin B** | SNIPER | 1.078 | 1.686 | 1.970 | 2.509 | 1.761 |
| | Rao et al. | 0.714 | 2.795 | 2.574 | 4.036 | 2.308 |

**Figure S8:** **(A)** SON TSA-seq and LaminB TSA-seq signal percentile distribution in SNIPER subcompartments in K562 (left) and the original subcompartments in GM12878 (Rao et al., 2014) (right). Distribution plots along the *x* and *y* axes represent the density of SON TSA-seq and LaminB TSA-seq signal, respectively, in each subcompartment. **(B)** Variance of SON TSA-seq and LaminB SON TSA-seq signal in each subcompartment in SNIPER K562 annotations and the original subcompartments in GM12878 (Rao et al., 2014).

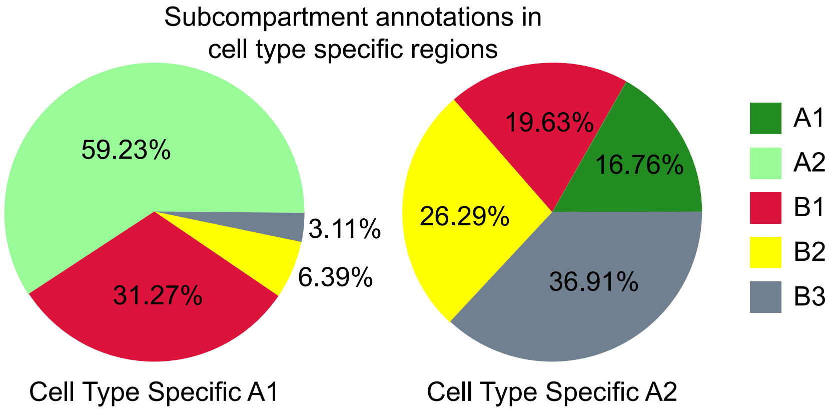


**Figure S9:** Fraction of subcompartment annotations at 100kb genomic regions in other cell types where A1 (left) and A2 (right) are specific to one cell type. In regions where A1 is specific to one cell type, the same regions in other cell types are frequently annotated as A2 (59.23%). In A2-specific regions, only 16.76% of regions in other cell types are in A1.

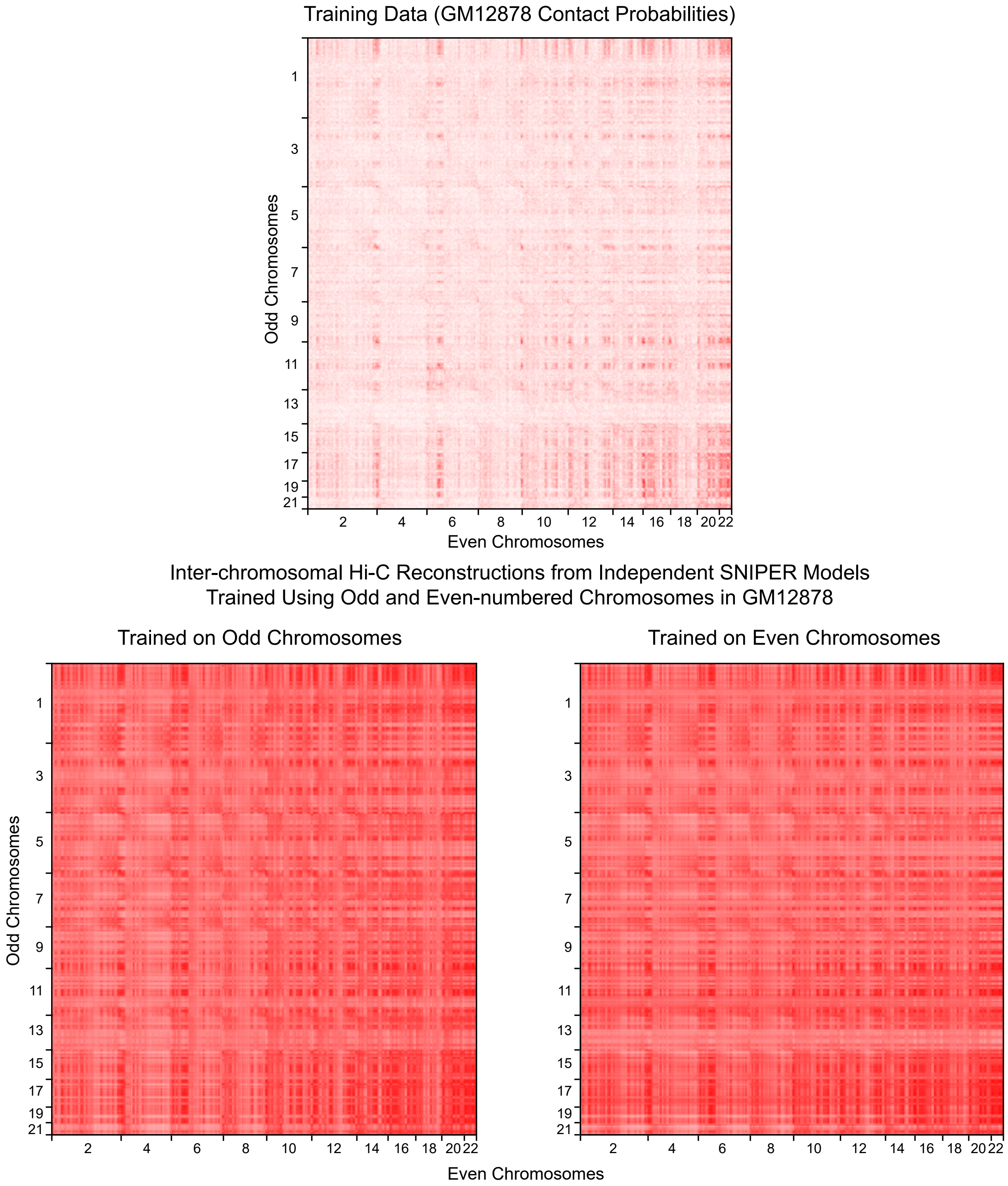


**Figure S10:** Hi-C probabilities reconstructed from sparse training data (top) using independent SNIPER models trained using odd (bottom-left) chromosomes and even (bottom-right) chromosomes.

A

**SNIPER Autoencoder 10-Fold Cross Validation (CE)**

| Fold | Training Loss | Test Loss |
|---|---|---|
| 1 | 0.632 | 0.636 |
| 2 | 0.630 | 0.630 |
| 3 | 0.632 | 0.633 |
| 4 | 0.631 | 0.634 |
| 5 | 0.631 | 0.633 |
| 6 | 0.632 | 0.632 |
| 7 | 0.631 | 0.633 |
| 8 | 0.630 | 0.631 |
| 9 | 0.633 | 0.634 |
| 10 | 0.634 | 0.635 |

B

**SNIPER Classifier 10-Fold Cross Validation Accuracy (%)**

| Fold | Overall | A1 | A2 | B1 | B2 | B3 |
|---|---|---|---|---|---|---|
| 1 | 91.97 | 90.50 | 94.83 | 84.80 | 90.00 | 94.93 |
| 2 | 91.02 | 94.65 | 91.89 | 87.08 | 86.89 | 91.53 |
| 3 | 92.60 | 98.25 | 91.73 | 82.69 | 88.75 | 96.71 |
| 4 | 92.44 | 95.92 | 95.19 | 87.69 | 80.42 | 94.96 |
| 5 | 91.65 | 95.30 | 89.53 | 86.17 | 82.35 | 97.24 |
| 6 | 92.36 | 97.95 | 87.45 | 87.94 | 89.68 | 95.35 |
| 7 | 92.44 | 96.75 | 88.93 | 89.25 | 89.60 | 94.83 |
| 8 | 91.81 | 96.46 | 90.78 | 84.13 | 88.98 | 94.25 |
| 9 | 92.43 | 97.15 | 90.48 | 87.17 | 86.76 | 95.08 |
| 10 | 92.75 | 97.55 | 90.73 | 84.13 | 90.84 | 96.02 |

**Table S1: (A)** 10-fold cross validation binary cross-entropy loss of the autoencoder in SNIPER. **(B)** 10-fold cross validation accuracy for each subcompartment in GM12878. Latent variable inputs were not balanced but each fold had similar accuracy to the classifier trained on the balanced training set.

**Prediction accuracy for different coverage levels**

| | Coverage (million Hi-C reads) | | | | |
|---|---|---|---|---|---|
| | 100 | 150 | 200 | 250 | 500 |
| Gaussian HMM | **47.88%** | **54.19%** | 56.10% | 77.30% | 75.27% |
| SNIPER | 37.61% | 49.71% | **63.55%** | **87.91%** | **90.83%** |

**Table S2:** Subcompartment annotation accuracy of the baseline Gaussian HMM and SNIPER at various levels of coverage. SNIPER outperforms the Gaussian HMM at 200 million read pairs and drastically improves prediction performance at 250 million read pairs.
